# Pattern Separation Contributes to Categorical Face Perception

**DOI:** 10.1101/2020.05.13.094276

**Authors:** Stevenson Baker, Ariana Youm, Yarden Levy, Morris Moscovitch, R. Shayna Rosenbaum

## Abstract

Traditionally considered a memory structure, the hippocampus has been shown to contribute to non-memory functions, from perception to language. Recent evidence suggests that the ability to differentiate highly confusable faces could involve pattern separation, a mnemonic process mediated by the hippocampal dentate gyrus (DG). Hippocampal involvement, however, may depend on existing face memories. To investigate these possibilities, we tested BL, a rare individual with bilateral lesions selective to the DG, and healthy controls. Both were administered morphed images of famous and nonfamous faces in a categorical perception (CP) identification and discrimination experiment. All participants exhibited nonlinear identification of famous faces with a midpoint category boundary. Controls identified newly learned nonfamous faces with lesser fidelity, while BL showed a notable shift in category boundary. When discriminating face pairs, controls showed typical CP effects of better between-category than within-category discrimination — but only for famous faces. BL showed extreme within-category “compression,” reflecting his tendency to pattern complete following suboptimal pattern separation. We provide the first evidence that pattern separation contributes to CP of faces.

---

A powerful image of Vladimir Putin and Donald Trump fused into a single face appeared on the July 30, 2018, edition of *Time* magazine (Figure 1). The image appears to have equal parts Trump and Putin. By straddling an identification boundary between the two world leaders, the illustration achieves the publication’s editorial purpose. The unsettling ambiguity created by the reduced discriminability of the hybrid image also highlights a unique property of human perception and memory: our natural bent to a) generalize categorically, and b) differentiate perceptually between items that lie along a physical (sensory) continuum. This process is known as categorical perception (CP)^1–4^. CP is clearly illustrated in our treatment of speech sounds, or colors in a rainbow, as discrete from one another, although they lie on uninterrupted continua of sound and light wavelengths, respectively^1,4^. The tendency to categorize and differentiate endures, even when a continuum is artificially created, as in the Putin-Trump mash-up^5–8^. Here, we examine the extent to which lesions to the dentate gyrus (DG) — a subfield of the hippocampus necessary for disambiguating similar visual input in memory through a process known as “pattern separation”^9–11^ — may also have differential effects on the perception of familiar and unfamiliar morphed faces.

**Figure 1.**
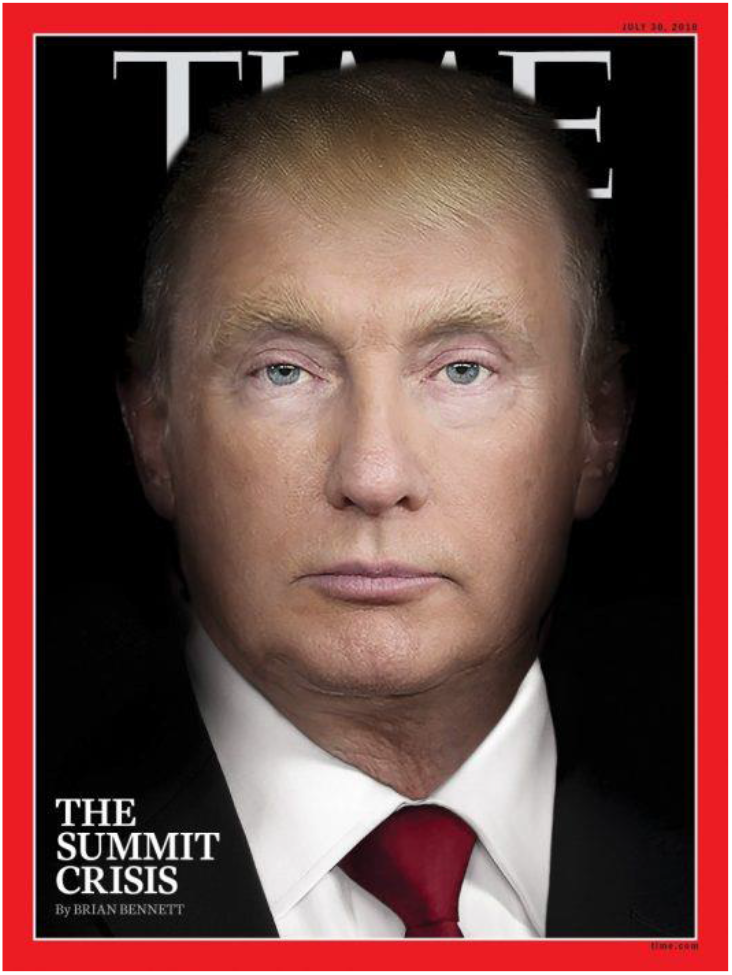
*TIME* Magazine, July 30, 2018

Although researchers have considered the core face network to be implicated in CP of faces^12^, none, to our knowledge, has suggested an essential role for the hippocampus or the DG. Indeed, most research on the discrimination of the feature-level representations of overlapping faces has found that, within the medial temporal lobe, this process recruits the perirhinal cortex, but not the hippocampus^13^. Recent evidence, however, hints that differentiating highly similar faces may be mediated by the hippocampus^14,15^, including through the process of pattern separation^16,17^. In addition, pattern separation within the DG has been speculated to facilitate the orthogonalization of overlapping face inputs^15^. It seems probable, furthermore, that face-name labeling, an essential part of CP testing of faces^5,7^, enlists associative encoding within the hippocampus, including in the DG, with decreasing activity in this subfield as face-name pairs are learned^18^. CP categorization or generalization mechanisms potentially also enlist the CA3 or CA1 hippocampal subfields, which receive inputs from the DG^11^. These areas are implicated in pattern completion/generalization functions of visual stimuli^10,19^ and could play a role in transforming face perceptual codes into memory representations^20^. Building upon the tentative evidence listed above, we believe that hippocampally mediated mnemonic processes subserve CP identification-discrimination paradigms, even those using high-interference face stimuli.

Complicating the question of whether pattern separation, completion, or generalization are factors in CP of faces, is the role played by personal expertise. Some believe that expertise and experience are crucial. For example, memory is often cited as a reason why CP effects seem stronger for familiar faces than for faces learned in the lab^5,7^. On the pattern separation side of the equation, perceptual expertise — as represented by a person’s pre-existing social categorization of faces, or the “other race” effect (ORE) — was found by Chang and colleagues^16^ to play a significant role in the ability of young adults to discriminate morphed faces. In this investigation, familiar same-race faces were more finely tuned (i.e., they had a sigmoidal or “S”-shape) as faces progressed across morphed steps from more-to-less similar^16^. Yaros et al. built upon these findings to propose a pattern separation model whereby successful behavioral discrimination across same-race faces is a function of the demands of memory relative to the degree of the overlap (i.e., interference) between face stimuli^17^.

Still, direct evidence of the interaction of CP with cognitive activities such as memory and learning, remains elusive^2,21^. Unanswered questions about the interaction of CP with higher-order processes are to be expected for a phenomenon manifested across modalities and involving top-down conceptual systems and bottom-up perceptual systems^3,21^. Nevertheless, researchers formulating neural network models of CP have reported several ways in which specific aspects of learning and memory are crucial components of the emergent properties responsible for transforming linear perceptual stimuli into nonlinear internal representations and subsequent categorical responses^1,21^. We hypothesized that the pattern separation function of the DG and the pattern completion/generalization functions of CA3/CA1, together with personal expertise, influence CP categorization/identification. Specifically, these elements push and pull on the operational markers of CP: the attendant effects of within-category “compression” (i.e., reducing perceived differences of faces within one identity category) and between-category “expansion” or “amplification” (i.e., expanding perceived differences of faces straddling either side of a category boundary)^3–5^.

In order to test this hypothesis, we presented BL, an amnesic person with a rare hippocampal lesion selective to the DG^22^, and healthy age-matched controls, with blended images of famous faces and nonfamous faces in a standard CP identification and discrimination experiment. We predicted that if CP relies on the mnemonic process of pattern separation, then BL would exhibit atypical behavior in identifying and discriminating nonfamous faces relative to famous ones, as the former would show the greatest reliance on the DG in learning new face-identity information. Meanwhile, CP tasks that rely on generalization (i.e., within-category discrimination) would be more dependent on pattern completion or generalization mediated, respectively, by BL’s intact CA3 and CA1^10,11,22^. The results of our tests provide a critical bridge between perception and memory in general, and CP and hippocampally mediated pattern separation in particular.

## Results

### Identification task

#### Control participants

Our experiment followed standard operationalization of CP and was divided into two phases^4^. During the identification phase, participants were asked to categorize images of faces. These faces were morphed along a continuum of contrasts between prototypes of two image identities. Participants specified, using a keystroke assigned to a name, whether the morphed face presented on the screen in front of them was more like one face or another. We expected nonlinear identification of the images. In other words, responses would show a sigmoidal or “S”-shaped change in identification as contrasts of the stimuli moved stepwise from one endpoint prototype (e.g., 90% of Ryan Gosling’s face and 10% of Benedict Cumberbatch’s face) to another (e.g., 10% of Ryan Gosling’s face and 90% of Benedict Cumberbatch’s face). The sigmoid activation function expresses the tendency of neurons to become “saturated” once a firing threshold (e.g., where perception changes qualitatively and confidently from one face to another) is reached^10^. We also expected that the sharpest change in classification would occur at the threshold (or category boundary). This boundary would be at, or near, a predicted point of subjective equality/maximum ambiguity (approximately 50%), where a concomitant change in identification from one face-name category to another typically occurs^5–8^.

Participant data for the two-interval forced-choice (2IFC) identification task of images of famous faces and nonfamous faces are summarized in Table 1 and graphed in Figure 2. We found the thresholds for famous faces (*M* = 49.75, *SE* = 0.75) and nonfamous faces (*M* = 50.48, *SE* = 0.66) to be close to the predicted point of subjective equality. We failed to find evidence that the gap between the two thresholds, −0.73, was significantly different, 95% CI [−2.65, 1.18], two-tailed *t*-test, *t*(34) = −.78, *p* = .44, *r* = .13. These findings illustrate the ability of middle-aged, healthy controls to categorize familiar and unfamiliar faces in a binary way. As a benchmark of CP, though, such an ability is a necessary but not a sufficient indicator of this phenomenon^23^.

**Figure 2.**
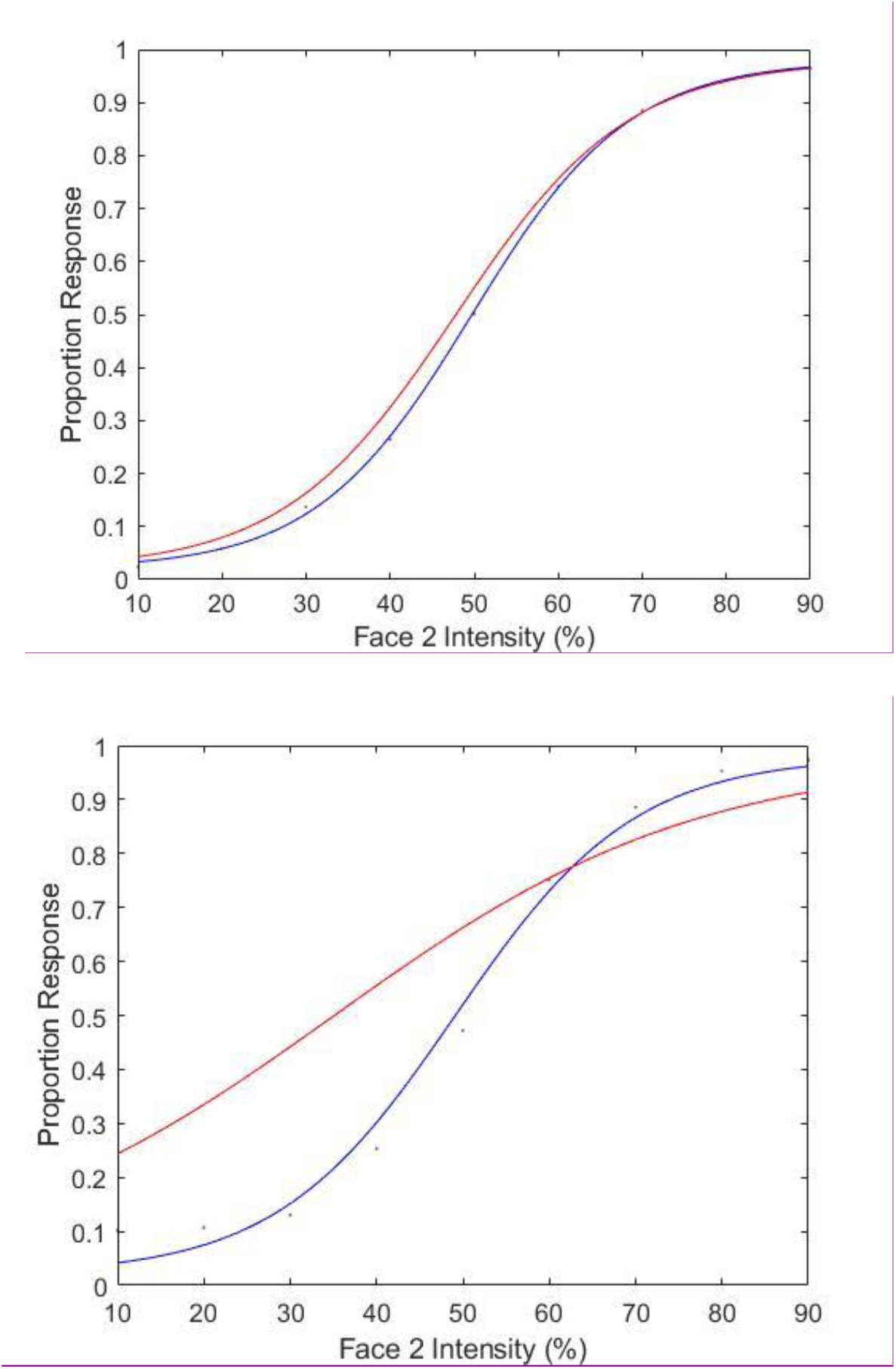
**Top:** Logistic function representing aggregate identification for controls for Famous Faces (blue) and BL (orange). **Bottom:** Logistic function representing aggregate identification for controls for Nonfamous Faces (blue) and BL (orange).

**Table 1.** Logistic Function Parameters.

| Group | Threshold ($\alpha$) | | Slope ($\beta$) | | Slope ($\kappa$) | |
| --- | --- | --- | --- | --- | --- | --- |
|  | FF | NF | FF | NF | FF | NF |
| Controls | 49.75 | 50.48 | 12.87 | 11.06 | 3.09 | 2.67 |
|  | (0.75) | (0.66) | (0.89) | (0.82) | (0.21) | (0.19) |
| BL | 47.82 | 35.12 | 9.76 | 4.74 | 2.34 | 1.14 |

*Note.* FF = famous faces. NF = nonfamous faces. Threshold ( $\alpha$ ) = category boundary (Face 2 intensity %). Slope ( $\beta$ ) = slope of the sigmoidal curve. Slope ( $\kappa$ ) = the slope of the curve at the contrast detection threshold (i.e., the first derivative of the slope). Values are averaged across participants in the group. Standard error appears within parentheses.

In accordance with previous CP studies using morphed faces^7,24^, we proceeded with an analysis of the threshold slopes. The difference, 0.42, in the steepness of the average slope at the contrast-detection threshold for famous faces (*M* = 3.09, *SE* = .21) and nonfamous faces (*M* = 2.67, *SE* = .19) did not reach significance, 95% CI [−.05, .89], two-tailed *t*-test, *t*(34) = 1.83, *p* = .076, although it did represent a medium-sized effect^25^, *r* = .30. In order to determine how well identification of famous faces and nonfamous faces aligned with respective sigmoidal curves, we ran a goodness-of-fit routine from the Palamedes toolbox for MATLAB^26^ on each participant’s data. We then analyzed the aggregate deviation scores in each condition using the chi-square cumulative distribution function, chi2cdf(x,v,’upper’), in MATLAB. We followed guidelines that an unacceptable fit corresponds to a probability value of *p* < 0.05^27^. Data for famous faces exceeded that value, chi2cdf (255.13, 245, ‘upper’), *p* = .32. By contrast, deviation scores for nonfamous faces did not reach the benchmark, chi2cdf (394.68, 245, ‘upper’), *p* <.001, indicating that their data points stray from the model fit^28^. An *a priori* assumption that the logistic model can best represent the identification of stimuli in a CP experiment was not met for both morphed face conditions.

#### Identification response times

As is illustrated in Figure 3, controls’ identification response times (RTs) for famous faces increased from the 10% endpoint of Face 2, across the 20%, 30%, and 40% morph steps to the peak point of ambiguity, 50%, then declined in a comparable stepwise fashion. Controls’ RTs for nonfamous faces revealed a more gradual rise and fall across the 40%, 50% and 60% morph steps. To investigate further whether controls classified famous faces and nonfamous faces differently, we compared identification response times between face endpoints (10% and 90%) with the peak point of ambiguity (50%). A 2×2 repeated-measures ANOVA revealed a main effect of endpoint *F*(1, 34) = 145.11, *p*<.001, *r* =.90, indicating that average endpoint response times (*M* = 1150.01, *SE* = 42.41) were significantly faster than the midpoint ones (*M* = 1577.01, *SE* = 62.50). A main effect of face familiarity, however, was not found, *F*(1, 34) = .067, *p*=.80, *r* = .04; nor did we detect a Familiarity × Face intensity interaction *F*(1, 34) = 1.70, *p* =.202, *r* = .22. This finding of slower RTs at, or near, boundary thresholds has also been documented using images of everyday objects^28^. As for faces, if people’s perceptions are equally torn between two competing identities, such as between the Trump and Putin face morph, then the time necessary to resolve this judgment will take longer than when the visual object is more obviously a “Trump” or a “Putin.”

**Figure 3.**
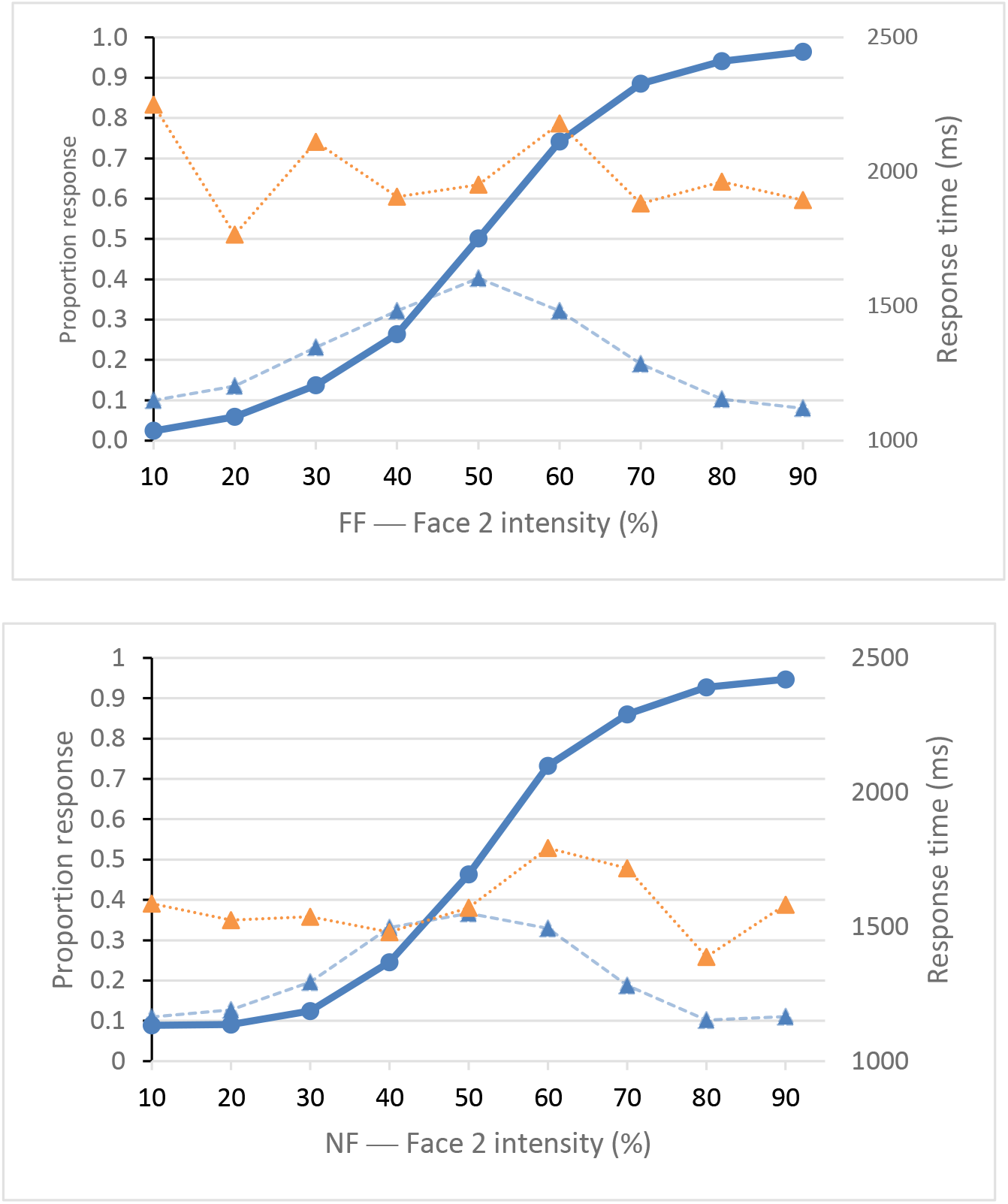
Identification Accuracy and Response Times. **Top**: Aggregate identification for controls for famous faces (FF, solid blue line), contrasted with response times for controls (dotted blue line) and BL (dotted orange). **Bottom** Aggregate identification for controls for nonfamous faces (NF, solid blue), contrasted with response times for controls (dotted blue) and BL (dotted orange).

BL’s RTs, however, failed to show this latency advantage for endpoint morphs. The Revised Standardized Difference Test^29^ revealed a significant difference between the average response times of BL and controls when contrasting endpoints (10% and 90%) and the threshold morph for famous faces (50%); *t*(34) = 3.31, *p* = .002. This difference reveals that BL did not exhibit the same speed advantage for identifying endpoint morphs over boundary morphs. A significant difference was also found when comparing nonfamous endpoints versus the boundary threshold (50% for controls and an average of 30% and 40% for BL); *t*(34) = 2.06, *p* = .047.

#### Threshold and slope values for BL vs. controls

Although results indicate BL had a contrast-detection threshold close to 50% for famous faces, his performance for nonfamous faces deviated from controls. In this latter condition, his category boundary was approximately 35% (Table 1 and Figure 2). In order to compare the category threshold of BL with the means of controls, we applied Crawford and Howell’s modified *t*-test for single cases^30^. Using this measure, we were unable to find a significant difference between BL and controls for the boundary for famous faces, *t*(34) = −0.43, *p* = .670. Indeed, BL’s results place him at approximately the 34th (i.e., average) percentile, 95% percentile CI [22, 47]. Additionally, the steepness of BL’s slope at this threshold was within normal limits, *t*(34) = −0.59, *p* = .281 (one-tailed), 95% percentile CI [17, 41]. Clearly, BL and controls showed categorical nonlinearity of response to a continuous linear change in the famous faces continuum^1^. In contrast, BL’s threshold for nonfamous faces was significantly different than that of controls, *t*(34) = −3.85, *p* < .001, .02 percentile, 95% percentile CI [0, 0.17]. We failed to find a significant difference for the steepness of BL’s slope at this boundary *t*(34) = −1.31, *p* = .20, 10th percentile, 95% percentile CI [4, 19]. The data indicate that, unlike controls, BL did not switch his responses from Face 1 to Face 2 near the middle of the continuum of noisy exemplars of unfamiliar faces, but did so earlier, around the 30% or 40% mark.

#### Predicted versus obtained discrimination (percent correct)

Table 2 shows average obtained discrimination accuracy (percent correct) for controls and BL for within- and between-category pairs. In order to determine how this performance compared with predicted accuracy (predicted from the classification task), we followed procedures for two-response classification detailed in Macmillan and Creelman (2005)^31^. This approach assumes that an observer’s discrimination response is unbiased and correlates with categorical labeling^4,31–33^. As can be seen in Table 3, predicted discrimination was estimated to be close to 51% (i.e., random) for within-category discrimination, and approximately 63% for cross-category pairs. These predictions correspond with the theoretical expectation that discrimination of morphed faces follows, and is actually influenced by, the nonlinear dynamics found in the identification of morphed faces: those faces that cannot be categorized differently will not be discriminated differently (as they are perceived as the same category/identity); those faces which can be categorized as distinct will also be discriminated as different (as they are perceived as a different category/identify)^2,4,33^.

**Table 2.** Obtained Discrimination Accuracy: Within- and Between-Categories (Percentage Correct)

| Group | FF within | FF between | NF within | NF between |
| --- | --- | --- | --- | --- |
| Controls | 35.54 (3.90) | 47.14 (4.77) | 43.21 (4.07) | 47.14 (4.35) |
| BL | 25.00 | 25.00 | 17.50 | 35.00 |

**Note.** FF = famous faces. NF = nonfamous faces. Controls and BL within = faces averaged across 10–30% and 70–90% morph steps; Controls between = faces in 40–60% morph step. BL FF between = faces in 40–60% morph step. BL NF between = faces in 20–40% morph step. Standard error provided in parentheses.

**Table 3.**
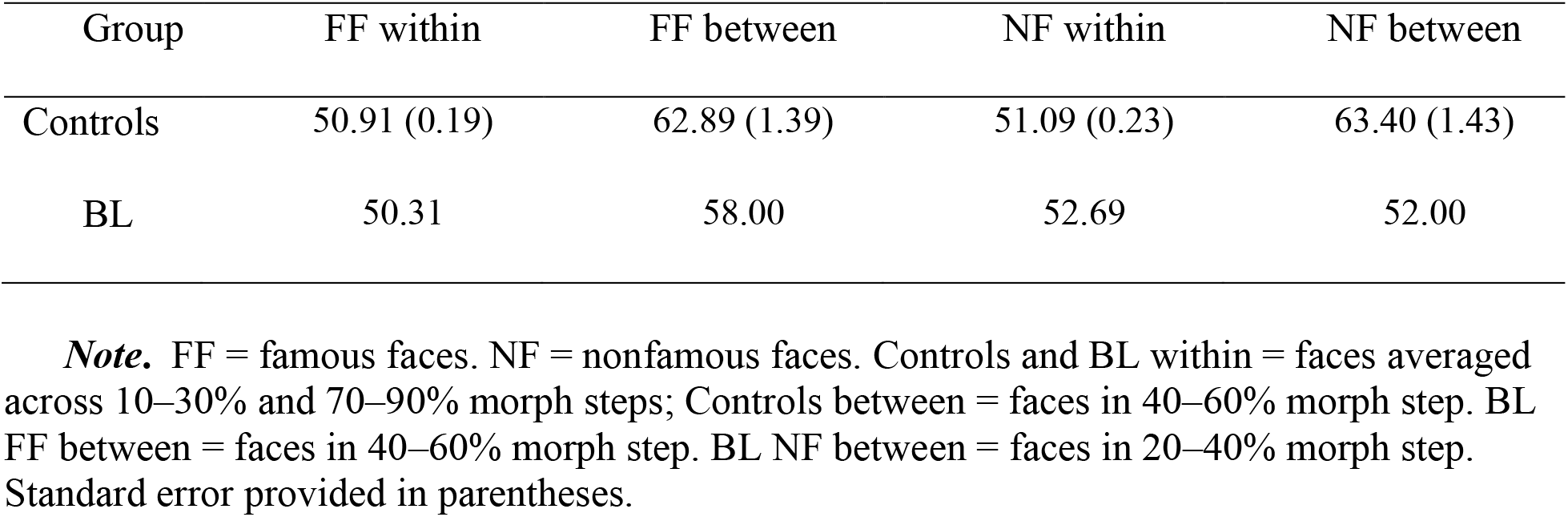
Predicted Discrimination Accuracy: Within and Between Categories (Percentage Correct)

Obtained discrimination accuracy (Table 2), presents a different picture. Controls were, 47.14% (famous faces) and 47.14% (nonfamous faces) accurate in between-category conditions and 35.54% (famous faces) and 43.21% (nonfamous faces) accurate in within-category conditions. BL achieved 25% discrimination accuracy in both famous face conditions. He was 35% accurate in the nonfamous between-category condition and 17.50% accurate in the nonfamous within-category condition. To compare obtained and predicted discrimination further, we compared controls’ average within- and between-category predictions versus obtained within- and between-category results. Paired *t*-tests, adjusted for multiple comparisons using the Holm-Bonferroni method^34^, revealed that the mean difference, −15.51%, between obtained accuracy (*M* = 41.46%, *SE* = 4.06%) and predicted accuracy (*M* = 56.97%, *SE* = 0.67%) for famous faces was significant, *t*(1,34) = −3.83, *p* = .001, 95% CI [−23.74%, −7.29%]. The mean difference, −11.80%, between obtained accuracy (*M* = 45.40%, *SE* = 3.98%) and predicted accuracy (*M* = 57.20%, *SE* = 0.70%) for nonfamous faces was also significant, *t*(1,34) = −3.03, *p* = .005, 95% CI [−19.72%, −3.90%]. Such subpar obtained relative to predicted discrimination was also observed in the performance of older adults (averaged across all morph steps) in a previous CP study of faces and attributed to the possibility that older adults do no use perceptual cues gained during the identification stage in later discrimination^7^. We believe that such low discrimination scores may also reflect the response bias frequently found in same-different discrimination paradigms^27,31^. Therefore, we next analyzed the discrimination data using *d*’ methods, which compensate for these forced-choice decision biases.

#### Obtained discrimination (*d*’) — controls

As the predicted versus obtained results help to illuminate the bias in participant responding evident in same-different tasks, a 2×2 (categorical boundary × familiarity) repeated-measures ANOVA was used to determine if control data varied across discrimination conditions. The dependent variable for this roving, forced-choice task, was discrimination *d*’ scores^27,31^. (See Table 4.) We found healthy controls performed significantly better in between-category trials compared to within-category trials, *F*(1,34) = 5.83, *p* = .021, *r* =.38. Face familiarity trended towards but did not reach significance *F*(1,34) = 3.33, *p* = .077, *r* =.30. The category boundary x familiarity interaction was significant *F*(1,34) = 8.20, *p* = .007, *r* =.44, prompting us to run an analysis of categorical boundary at each level of face familiarity. Paired *t*-tests, adjusted for multiple comparisons using the Holm-Bonferroni method^34^, showed that within-category famous faces (*M* = 1.74, *SE* = 0.16) were discriminated at a lower rate than between-category famous faces (*M* = 2.23, *SE* = 0.18). This difference, −0.49, 95% CI [−.75, −.23] was significant, *t*(34) = −3.84, *p* = .001, *r* = .55. However, there was no indication that within-category nonfamous faces had a similar perceptual disadvantage relative to between-category ones, *t*(1,34) = .002, *p* = .998. The latter result is unsurprising, as the within- and between-category *d*’ values for nonfamous faces were nearly the same. Moreover, within-category famous faces were discriminated at a lower rate than within-category nonfamous faces (*M* = 2.13, *SE* = 0.18). This difference, −0.39, 95% CI [−.61, −.17] was significant, *t*(34) = −3.63, *p* = .001, *r* = .53. A similar finding was not evident when comparing between-category famous faces with between-category nonfamous faces, *t*(1,34) = 0.79, *p* = .437, *r* = .13.

**Table 4.** Discrimination Accuracy: Within- and Between-Categories (d’)

| Group | FF | FF | NF | NF |
| --- | --- | --- | --- | --- |
|  | within | between | within | between |
| Controls | 1.74<br>(0.16) | 2.23<br>(0.18) | 2.13<br>(0.18) | 2.13<br>(0.18) |
| BL (all face pairs) | 0.43 | 1.48 | 0.00 | 1.16 |
| *t* | −1.35 | −0.71 | −2.00 | −0.91 |
| *P (one-tailed)* | 0.093 | 0.243 | 0.027 | 0.186 |
| BL (identifiable faces) | 0.60 | 2.08 | 0.00 | 1.59 |
| *t* | −1.17 | −0.14 | −2.00 | −0.51 |
| *P (one-tailed)* | 0.124 | 0.444 | 0.027 | 0.308 |
| BL (unidentifiable faces) | 0.00 | 1.16 | 0.00 | .53 |
| *t* | −1.79 | −1.01 | −2.00 | −1.50 |
| *P (one-tailed)* | 0.041 | 0.161 | 0.027 | 0.072 |

**Note.** FF = famous faces. NF = nonfamous faces. Controls and BL within = faces averaged across 10–30% and 70–90% morph steps; Controls between = faces in 40–60% morph step. BL FF between = faces in 40–60% morph step. BL NF between = faces in 20–40% morph step. Standard error provided in parentheses. Values of Crawford *t*-test comparisons between patients and controls are provided in *t* (34) and *p* (one-tailed).

#### Obtained discrimination (*d*’) — BL

Discrimination accuracy rates for BL are reported in Table 4. As can be seen here, his lowest score was found in within-category nonfamous faces. BL had a *d*’ score of 0 (i.e., random responding) in this condition. Using Crawford and Howell’s modified *t*-test for single cases^30^, we found this difference between BL and controls for nonfamous faces, −2.13, was significant, *t*(34) = −2.00, *p* = 0.027, one-tailed. BL’s lack of sensitivity in this condition was apparent even when we only included those face-pairs that BL could identify by name in a post-test (see Table 4 and Figure 4). A significant difference between BL and controls was also found when comparing the within-category famous faces for which he could not assign a person-identity label *t*(1,34) = −1.79, *p* = .041, one-tailed.

**Figure 4.**
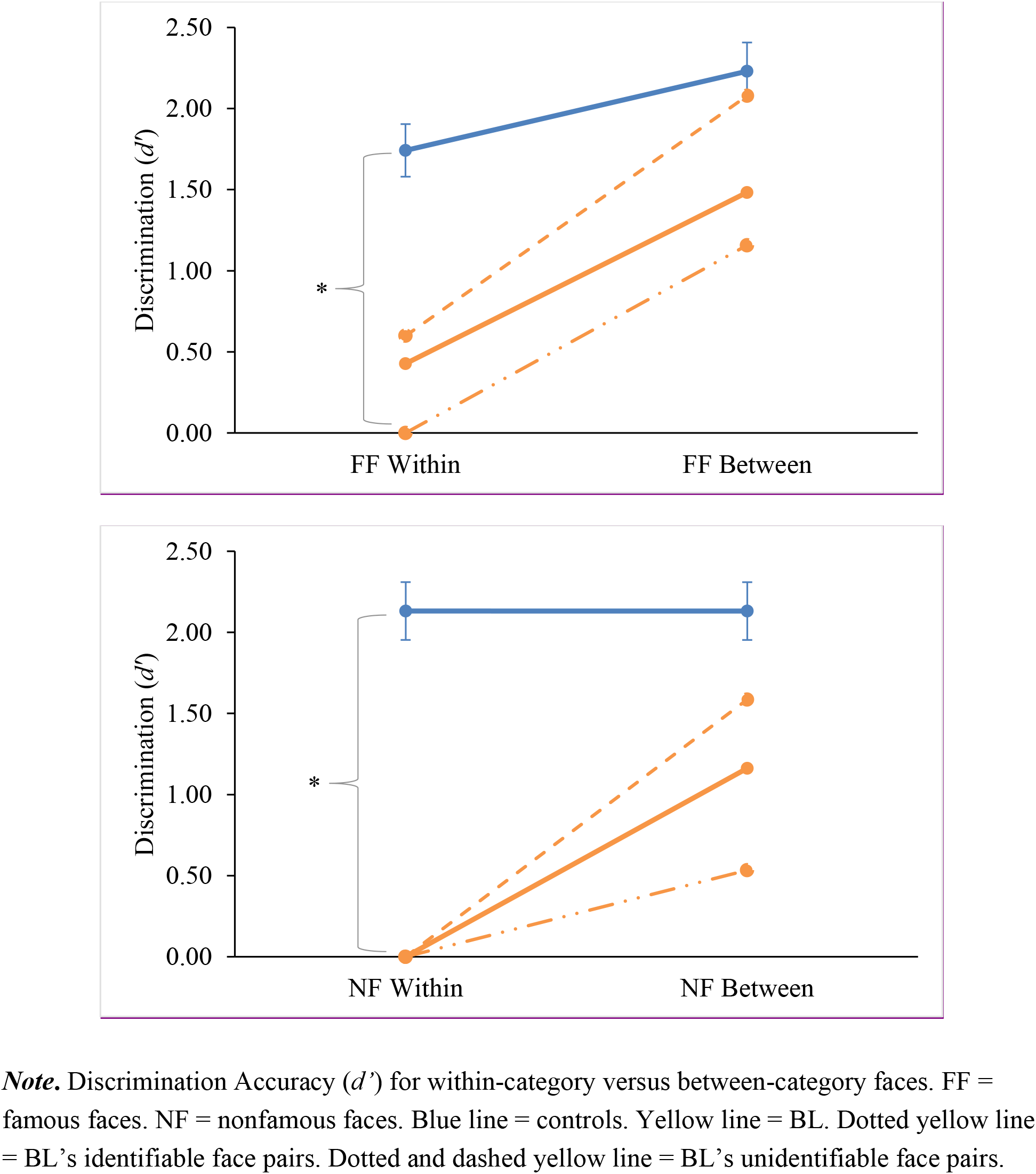

## Discussion

We set out to determine if CP, like pattern separation, is modulated by hippocampal DG integrity and is influenced by semantic familiarity. As predicted, healthy controls exhibited CP effects for famous morphs — as defined by successful categorical identification alongside within-category compression and between-category expansion^3^. On the other hand, CP effects were absent for nonfamous morphs. These data suggest that personal familiarity influences the identification and discrimination of highly confusable face images. In contrast, BL, an individual with selective bilateral ischemic lesions of the DG, exhibited idiosyncratic identification of nonfamous faces and significantly lower accuracy rates for within-category discrimination of both nonfamous and famous faces. These indicators implicate DG integrity in supporting the discrimination of similar faces. Furthermore, BL did not show significantly faster identification response times for endpoint faces relative to boundary ones, suggesting that newly learned information about morphed faces may not have been pattern separated to the degree necessary to influence efficient decision-making. Together, these findings indicate that the perceptual act of CP and the mnemonic act of pattern separation are interrelated through underlying cognitive processes via a common hippocampal substrate.

Similar perceptual and mnemonic relationships can be pieced together from process, psychophysics, or neuroanatomical approaches to CP or pattern separation. In a process approach, such interpretation is conceptualized as the tendency for individuals to categorize and discriminate based on stored information, such as category exemplars or prototypes^1,21^ Indeed, early neural network models of CP incorporated mechanisms of associative memory based upon prototype models. According to these models, noisy perceptual stimuli are replaced by noise-free prototypes from memory^1,21^. Within the scope of such approaches, reliance on previously stored exemplar information (which depends on semantic and episodic memory), would necessitate the ability to maintain the distinctiveness of this information through hippocampal pattern separation^35^. Retrieving a canonical pattern to replace a noisy one would also, presumably, engage the autoassociative circuitry of CA3^20^. Pattern separation activity within the DG/CA3, therefore, would track or support CP.

The interrelatedness of CP and pattern separation also follows from the parallels in the conclusions that can be drawn from the experimental paradigms used to operationalize the phenomena. For example, the CP task begins with the repeated presentation of morphed faces. In theory, some of these faces with overlapping features (at least the more recognizable ones nearest the endpoints) are sparsely coded by pattern separation processes^17^. The orthogonal episodic representations of each morphed face are then projected from the DG onto the CA3 hippocampal cell layer via the mossy fiber pathway^10,11^. Recurrent collaterals within the CA3 ensure the episodic elements of these learning episodes are bound together and are completable from partial cues^10,11^.

In this manner, the CP task resembles a widely used test of pattern separation, the Mnemonic Similarity Task (MST)^36^. Yet the MST saves its highly similar stimuli, or lures, for the test phase. During the study phase of the MST, the everyday visual objects presented to participants (e.g., saxophones, sea horses, picnic baskets) are more dissimilar than the famous and nonfamous faces used in our task. Moreover, the CP task does not involve a recognition memory test phase, as does the MST. Nevertheless, the morphed faces in the CP paradigm would be pattern separated regardless of whether they were later tested in memory; in effect, the CP discrimination phase is an assessment of how well the faces are pattern separated. Due to his DG lesion, BL was unable to pattern separate highly similar nonfamous faces as well as controls during the identification phase. As a result, he lacked the sparse encoding necessary for perceptual, fine-tuned discrimination.

Based on BL’s memory performance on the MST in a previous study, where he was found to be able to recognize highly dissimilar foils^22^, we surmise that he did have the ability to encode coarser face representations. These more generalized representations likely prompted him to autoassociate highly similar within-category faces to an endpoint identity. This interpretation could explain why BL’s discrimination for within-category nonfamous faces was at floor and significantly lower than that of controls. To BL, all within-category faces were indistinguishable (i.e., they were perceived absolutely), a CP “ideal case”^4^ rarely achieved through experimentation^2,4^. BL’s slightly improved performance for those between-category faces he could name in a post-test could reflect the fact that faces with more meaningful associations can be supported by extra-hippocampal, conceptual structures^37^. These may have helped to shift the representation of faces from being perceptually image-based to being conceptually view-invariant^38^. Consequently, the faces that he could name using person-identity information offered less interference, facilitating discrimination in between-category decision-making, the same way that pattern separation of morphed faces varies depending upon the relative familiarity and invariance of the faces involved in any particular trial^16,17^.

When presented with famous faces, BL’s ability to discriminate within-category morphs was also at or near floor. The famous faces he could not name in a post-test, like nonfamous faces, were discriminated at significantly lower rates than those correctly named by controls. Controls, meanwhile, were worse at identifying within-category famous faces than between-category ones, revealing one of the definitional criteria of CP effects^3,4^ (albeit without BL’s apparent total within-category indiscriminability). This within-category compression shown by BL and controls may reflect something akin to a “perceptual magnet effect”^39^. We believe this perceptual pull is caused by shared connectivity within CA3. It works to elicit “same,” rather than “different,” judgments, leading to reduced *d*’ scores for highly similar within-category images, consistent with CA3’s role in pattern completion.

As we demonstrated through an analysis of identification response times, healthy controls showed slower responses for face morphs at the maximum point of ambiguity (50%) compared with endpoint faces (10% and 90%). One explanation for this finding is that these data represent the influence of attractor networks, as mental representations compete against each other for resolution within CA3 attractor basins^10,28,40^. The peak RTs correspond to those perceptual decisions which take the longest to resolve. In all likelihood, these decisions rely on the ability to see the prototype masked within the morph, an ability which recruits face memory recognition processes within the dentate gyrus of the hippocampus^14,16,17^.

Overall, our findings indicate that the role of the hippocampus in face identification and discrimination needs to be taken into account along with the roles of structures within the core face network, such as the lateral occipital cortex, fusiform gyrus, anterior temporal cortex, and prefrontal cortex^41–44^. Having established that the hippocampus contributes to at least one aspect of face perception^14,15,45,46^, we hoped to determine how it interacts with other regions of the core face network. Thus, the name of the face, a person’s familiarity with the face, and the myriad of meaningful associations and emotional biases or connections humans have for depictions of other humans, such as in the Trump-Putin morph, are equally important. These elements bring the conceptual richness of any face into being, as described in classic cognitive models of face processing^51^. For any familiar or newly learned face, the semantic face-name identity is particularly important. The ability to disambiguate the nose and eyes of Ryan Gosling from the chin and cheekbones of Benedict Cumberbatch should have little bearing on consciousness if one could not assign a holistic identity to these features. This identity information is associated with faces by the hippocampus and neocortex^18,48^, and, we believe, is central to the identification and discrimination of highly similar faces.

A growing literature suggests that the MTL may engage in different expressions of mnemonic or perceptual processing, possibly as a result of its projections to and from the neocortex^49^. Previously, the perirhinal cortex was thought to be the last stop within the MTL for this process, with perceptual contributions of the hippocampus limited to the spatial domain^13^ or to relational processing^46^. Using a previously established measure of CP for faces, we were able to draw a parallel between both phases of identification and discrimination for controls and for a person with a focal brain lesion and a deficit in behavioral pattern separation. In doing so, our findings provide strong direct evidence that the hippocampus, and the DG in particular, also aid in processing faces in a high-interference task. These abilities are necessary for CP. Insofar as CP has been shown to be functionally dependent on perceiving differences and similarities in perceptual data, it appears to be intertwined with mnemonic pattern separation. The current study brings us closer to understanding this relationship and, more generally, how the hippocampus enables both perception and memory.

## Methods

We followed a standard CP identification and discrimination approach^4,50^ while expanding upon previous studies in several ways. First, during an initial identification phase, we applied a goodness-of-fit measure to examine the application of the traditional logistic psychometric function used to model classification. This function relates the proportion of trials assigned to a progressive series of identification steps to the intensity of the stimulus in that continuum. When graphed, the function takes on a familiar sigmoidal or “S”-shaped curve. Second, we analyzed the discrimination data using signal detection measures, thought to be the best practice for same-different tasks^5,31^. Third, we acknowledged that finding a model fit with distinct labeling categories and a sharp boundary for the data during identification is not in itself a sufficient benchmark for CP^23^. Instead, it is the first of four operational criteria traditionally used to demonstrate CP^4,51^. A second criterion — being able to predict the discrimination function from the identification function — was not found in our experiment, although this hypothesis is often unfulfilled in CP studies^4,31^.

The third and fourth criteria are tested during the discrimination phase^33^. Here, participants in CP experiments must show a detectable advantage, or “peaks”,^3,4^ in discriminating stimuli that cross a category boundary (e.g., 40% of Benedict Cumberbatch’s face and 60% of Benedict Cumberbatch’s face). A closely related fourth criterion is the presence of “troughs” of inferior discrimination^3,4^ for within-category stimuli (e.g., 70% of Benedict Cumberbatch’s face and 90% of Benedict Cumberbatch’s face). Alternately, some researchers have pointed out that the missing ingredient in CP studies is more often the failure to find chance discrimination for within-category discrimination; when that occurs, it has been suggested that other terms, such as “category boundary effect,” be used in place of CP^51^.

### Participants

#### Control participants

CP was assessed in 37 neurotypical adults, matched in age and education to an individual with bilateral DG lesions and impaired pattern separation (BL, described next). Two controls were excluded from final analyses for having discrimination/identification scores greater than 2.5 standard deviations below mean scores, resulting in a final test group of 35 control participants (20 females, *M*_age_ = 56 years, age range: 50 − 64 years). All controls were recruited from the community via advertisements or from participant databases at York University and Baycrest Health Sciences, and all had a Montreal Cognitive Assessment (MoCA) score at or above 26 (out of a 30-point maximum). Participants provided written, informed consent in accordance with the ethics review boards at York University and Baycrest and to the standards of the Canadian Tri-Council Research Ethics guidelines.

#### BL

To better understand how the DG region is also involved in learning and representing face categories, we examined CP abilities in BL, a memory-impaired individual with lesions to the DG. BL, who was 56 years old at the time of testing, sustained a hypoxic-ischemic brain injury in 1985 following an electrical injury and cardiac arrest^52^. In 2015, high-resolution 3T MRI scans of BL’s hippocampus revealed bilateral ischemic lesions that appeared to be restricted to the DG and a portion of the CA3 hippocampal subfield^22,52^. Neuropsychological assessment of BL revealed mild anterograde amnesia and moderate retrograde amnesia; facial recognition, as measured by the Benton Facial Recognition Test, was recently found to be within normal limits. To ensure reliability in performance in a single case, BL was tested on three different occasions (September 2017, December 2017, and October 2018). Only results of the last session are reported here, as it was the only one in which the experimenter responded on behalf of BL, in order to reduce the cognitive load placed upon him during task performance, and it was also the only testing session which incorporated a post-test of his face-identity knowledge.

### Face Stimuli

Evidence from Beale and Keil (1995)^6^ have suggested that categorical perception can occur when using faces as a categorical item and morphing one face to another along an artificial continuum. For the present study, stimuli selection was initiated by downloading pictures from the Internet of faces of American and Canadian public figures (e.g., politicians, actors, musicians) who became famous over the last 30 years. In addition, pictures of faces of people who were not famous or recognizable were selected from an existing database^13^. We then paired faces according to age, race, and gender (e.g., Ryan Gosling and Benedict Cumberbatch).

All faces/face pairs were then processed according to Lee et al. (2014)^7^. Briefly, faces were cropped into ovals. The resulting cropped images were matched in pairs, and the face pairs were morphed using a facial morphing program at nine 10-degree intervals and labeled for the percentage of Face 2 in the pair. For example, 90% Benedict contained 10% Ryan and 90% Benedict; 80% Benedict contained 20% Ryan and 80% Benedict; 70% Benedict contained 30% Ryan and 70% Benedict; and, 60% Benedict contained 40% Ryan and 60% Benedict. In addition, an equal morph pair was created (50% Ryan and 50% Benedict). See Figure 5.

**Figure 5.**
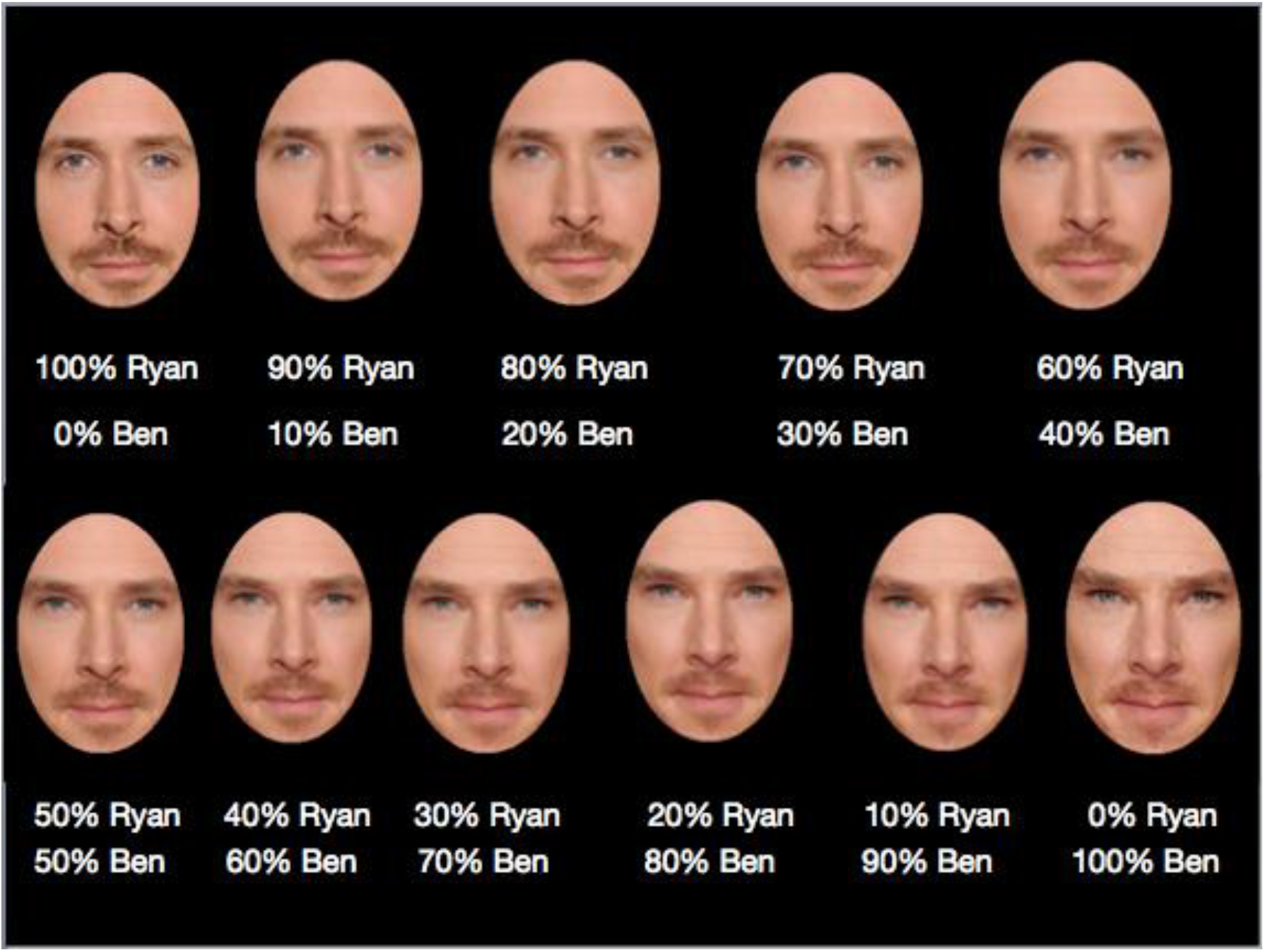
Example of exemplar morphs of famous faces (Ryan Gosling and Benedict Cumberbatch).

Each participant was tested with a Dell Latitude E5540 computer (Intel Core i7-4600U). The monitor was 15.5 inches with a 1366 × 768 × 60-hertz resolution. The experimental stimuli were presented with E-Prime 2.0.

### Procedure

#### Famous face recognition task

Before testing, participants completed a famous face recognition task, allowing the experimenter to select a group of famous faces known to the participant. During this test, the experimenter read through a list of famous names, and the participant was required to note the context in which the celebrity was famous. Additionally, the participant was required to describe the general appearance of the celebrity. Once the participant was able to recognize a list of at least four famous face pairs, they proceeded to the classification task.

#### Classification task

The classification task followed procedures set out by Lee et al. (2014)^7^.These procedures are briefly reported here. A training phase preceded the classification task. Participants were presented with cropped face pairs shown side-by-side and without morphing (e.g., 100% Ryan and 100% Ben). Below each face, the real name of the celebrity, or an invented name for the nonfamous person, was displayed. Participants had two minutes to study the pair and the affiliated names. The training phase was accompanied by a recognition test of single faces (at 100%) to ensure participants could label each face by its appropriate name (two choices were given). Participants could only proceed to the classification task after successfully labeling the famous and nonfamous faces to be used in testing.

During the classification task, participants were presented, one at a time, with four pairs of famous faces and four pairs of nonfamous faces. As described above, there were nine morphed image levels of each Face1-Face 2 contrast (named for the intensity level of Face 2): 10%, 20%, 30%, 40%, 50%, 60%, 70%, 80%, 90%. Each morphed image trial consisted of one face presented in the middle of the screen, with one name appearing above the face and one name presented below the face. These trials were randomly presented five times per morph pair per morph step, resulting in each Face1–Face2 contrast being presented 45 times across the different levels of morphing (180 trials in total for each face condition). Participants were instructed to respond to the identification question, *Who is this?*, for each morph pair by clicking the up or down arrow key in the direction of the perceived identity name.

#### Same-Different Discrimination task

As with the classification task, the discrimination task followed procedures set out by Lee et al. (2014)^7^, briefly reported here. The discrimination task began with participants studying each of the four famous and four nonfamous face pairs at 100%. Afterwards, the participants were randomly presented with 62 trials per pair at the following intensity levels of Face 2: 10-10%; 10-30%; 20-20%, 20-40%, 30-30%, 30-50%, 40-40%, 40-60%, 50-50%, 50-70%, 60-60%, 60-80%, 70-70%, 70-90%, 80-80%, 90-90%. The images presented in each trial were identical to those presented during the identification task. Participants were instructed to focus on the two images presented in each trial as opposed to the identity of the faces and to respond by way of an arrow press if the images presented were the *same* or *different*.

#### Scene classification and discrimination task

At the same time, we administered the experiment with famous and nonfamous faces, we also ran a pilot identification and discrimination task. It consisted of morphed scene stimuli. It was run on controls and patient BL (during his first two testing sessions). The scene trials were ordered after the face trials (i.e., faces classification, scene classification, faces discrimination, scene discrimination). This data is unreported here.

### Analyses

#### Identification

Endpoint accuracy, or the ability of participants to learn and identify faces at 90% intensity of Face 1 or Face 2, was first evaluated across conditions. Controls were found to be 92-97% accurate in correctly identifying the famous and nonfamous endpoint faces. Subsequent analyses of the identification data was done using SPSS Version 26 or the Palamedes toolbox^26^ for MATLAB. The category boundary dividing categorical perception of Face 1 and Face 2 was inferred through finding the slope of the logistic function at its contrast-detection threshold, assumed to be the steepest point of the sigmoidal curve^7,24,26^. The threshold is the point at which the correct proportion of responses reaches a criterion, typically 50% in CP experiments using faces^5–8,53^. This point aligns with the category boundary and is the approximate threshold at which participants began to identify a morphed face as Face 2, more than as Face 1. The model fit of the data, as defined by the parameters of the logistic function (Table 1) was determined through routines to evaluate psychometric functions selected from the Palamedes toolbox^26^ for MATLAB.

#### Discrimination

In order to use signal detection measures^31^ for evaluating the unbiased discrimination sensitivity of participants, we first averaged hits (percentage correct) for endpoint morph steps (10–30% and 70–90%). These pairs represented within-category faces and were found to be strongly correlated and without statistical variation in both face conditions. We also calculated hits for between-category faces, using the 40–60% morph step. False alarms were calculated for the respective identical pairs which could be matched with the within-or between-category endpoints (e.g., 10–10%, 60–60%). False alarms were calculated as the ratio of incorrectly responding *different* to these same pair trials. The *d*’ values for within-category faces and between-category faces were then determined using Palamedes MATLAB routines for a forced-choice same-different task within a differencing model (one in which stimuli rove along a continuum)^27,31^. The decision strategy assumed by this model is that participants respond *different* only when the perceived difference in each pair exceeds a criterion^27,31^. This criterion is thought to be conservative in same-different tasks, with observers more likely to answer “same” than different. The *d*’ strategy outlined above responds to this bias inherent in same-different tasks^26,27,31^. Subsequent analysis of the discrimination data in SPSS Version 26 took advantage of the factorial design of the experiment to produce a 2×2 ANOVA. The factors were Familiarity (two levels) and Categorical Boundary (two levels), with the dependent variable being discrimination *d*’.

#### Patient comparison

To ensure the reliability of findings in single cases, we used Crawford and Howell’s modified *t*-test for single cases^30^. This test treats the control sample’s data as statistics, rather than as parameters, controlling for Type I errors when testing whether a single case’s score is significantly below that of controls^30^. It also provides estimates of the percentage of the normal population falling below a single case’s score, as well as the confidence interval (CI) on the observed result. Additionally, the difference between BL and controls on two tasks was examined using Crawford & Garthwaite’s Revised Standardized Difference Test (RSDT) ^29^.

## Acknowledgements

This study was funded by a Vision: Science to Applications (VISTA) York Research Chair and NSERC Grant RGPIN-04238-2015 to RSR and NSERC Grant A837 to MM. We thank Dr. Nicolaas Prins for assistance with statistical analyses. We also acknowledge Dr. Yunjo Lee, as well as Nick Hoang and Marilyn Ziegler for helping to implement stimuli developed in their lab.

## Author contributions

S.B., A.Y., M.M. and R.S.R. conceived the project and designed the experiment. Data collection was performed by S.B., A.Y. and Y.L. Data analysis was performed by S.B., A.Y., Y.L., M.M. and R.S.R. The paper was drafted by S.B. M.M., and R.S.R., and AY and Y.L. provided critical input on revisions. All authors approved the final version of the paper for submission.

## Data availability

The data that support the findings of this study are available on request from the corresponding authors (S.B. or R.S.R.).

